# OmegaSwitch: Bayesian Markov-Modulated Codon Models for Estimating *dN/dS*

**DOI:** 10.64898/2026.08.14.744968

**Authors:** Wesley C. DeMontigny, Charles F. Delwiche

## Abstract

Selective pressures can vary across both sites and evolutionary lineages; however, most codon models accommodate heterogeneity along only one of these dimensions and require the number of selective regimes to be specified in advance. Here, we introduce *OmegaSwitch*, a Bayesian phylogenetic software framework for inferring changes in the nonsynonymous-to-synonymous substitution-rate ratio (*dN/dS*) across sites and through evolutionary time. We implement a Markov-modulated codon model in which lineages transition among discrete *dN/dS* regimes and use reversible-jump Markov chain Monte Carlo to infer the number of regimes simultaneously. We further develop a Dirichlet-process mixture extension that allows the parameters governing these time-heterogeneous processes to vary among sites. Ancestral sampling produces joint posterior distributions of *dN/dS* across sites and nodes of the phylogeny, enabling lineage- and site-specific summaries with quantified uncertainty. Simulation analyses showed that both the posterior intervals for *dN/dS* and the number of evolutionary regimes were well calibrated under both models. We demonstrate *OmegaSwitch* using vertebrate *α*- and *β*-globins. *OmegaSwitch* therefore provides a flexible Bayesian framework for investigating how selective pressures vary across protein-coding sequences and phylogenetic history.

## Introduction

Selective pressures are a central concept in evolutionary biology. They can be defined as factors that favor the reproduction of organisms with particular traits. Modern evolutionary biologists are commonly interested in the strength and nature of selection, and often those acting on specific sites in a protein-coding gene. Rather than directly mutating amino acid residues and observing the effect on fitness, we use inferred rates of synonymous and nonsynonymous substitutions to indicate how selection is acting; we summarize these rates as the ratio of non-synonymous to synonymous substitution rates, called the *dN/dS* ratio (1). Highly conserved sites have a low *dN/dS* ratio; when conservation is observed across a wide range of taxa in a clade of interest, it is inferred that members of the clade with non-synonymous mutations are undetected because lineages with these mutations die out. Such conserved sites are said to be under negative selection. Likewise, when a site is highly variable, it will have a high *dN/dS* ratio; when this pattern is observed across the entire dataset, it is inferred that innovation at the site has been beneficial for fitness or that variation at the site in question is of little consequence. These variable sites are said to be undergoing positive selection.

Codon models are the standard way for estimating *dN/dS* ratios. The simplest codon models that measure the *dN/dS* ratio characterize the protein as a whole; these models are unable to identify the selection regimes of particular sites, making their measured *dN/dS* not particularly useful, as it is unsurprising that proteins, on average, are undergoing negative selection (2). Additionally, selection varies across sites and throughout time. Not all sites in a protein are equally constrained structurally and functionally (3), and processes such as gene duplication or gain-of-function substantially alter how a gene family evolves along a phylogenetic tree (4–6). Therefore, these simple codon models have since been extended to detect varying selection at particular sites, branches, or both. Frequentist implementations of these models commonly employ likelihood-ratio tests between nested hypotheses (1, 7–9), which can be powerful and computationally efficient, but are often constrained in the questions they answer. Branch and branch-site tests, for instance, detect positive selection only on “foreground” branches designated by the user, assuming neutrality or negative selection elsewhere (7). This can make interpretation challenging when selection may have acted more broadly (10). Fully Bayesian approaches address some of these limitations by modeling the entire generative process, producing posterior distributions over *dN/dS* parameters and related metrics, along with their associated uncertainty. This ultimately supports a broader range of questions about the evolutionary process. For example, posterior samples can be used to quantify differences in *dN/dS* among sites, taxa, or ancestral lineages, or to calculate posterior probabilities for arbitrary comparisons without fitting an additional model.

We draw inspiration from two probabilistic models that allow selection to vary across sites and over time. First, Huelsenbeck et al. (11) developed a Bayesian non-parametric model using a Chinese restaurant process (CRP) to model site-specific *dN/dS* ratios. The Bayesian non-parametric approach is particularly attractive because it allows the model to scale in complexity based on what the data suggest, rather than specifying a fixed number of selection categories. Using this model, they showed heterogeneity in the selection acting on each site of various viral protein-encoding genes. Second, Guindon et al. (12) proposed a series of Markov-modulated models that allow the evolutionary model to vary over time and across sites. The parameters of these models were inferred using maximum likelihood, and they utilized frequentist model comparison to select the optimal number of selective regimes. They were particularly interested in the expected frequency with which a selective regime would appear on a branch, which could indicate bursts of increased selection. However, they did not leverage the fact that their Markov-modulated models could be simulated to produce ancestral *dN/dS* states and to identify the regime likely to be active.

Here, we implement models from (12) in a Bayesian setting and use reversible-jump Markov chain Monte Carlo to perform automatic model selection and inference of *dN/dS* heterogeneity across time and sites. Second, we extend (11) to use an infinite mixture of Markov-modulated rate matrices, allowing time-heterogeneous processes to vary across sites. Under these models, we perform ancestral sampling to infer a joint posterior of *dN/dS* at each element of an MSA (site and taxon). We develop these models in a novel software, *OmegaSwitch*, and analyze the coverage of the model’s posterior credibility intervals through simulation analyses. To provide a well-understood demonstration of our software, we analyze selection on vertebrate *α*- and *β*-globin data using both models. We demonstrate the flexibility of our models in producing useful posterior statistics.

## Materials and Methods

### Markov-Modulated Codon Models

Codon models assume that substitutions on a phylogenetic tree occur according to a continuous-time Markov chain with generator matrix (13):

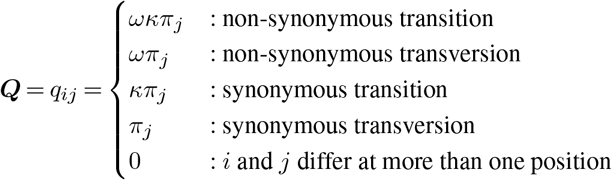

where *ω* is the non-synonymous rate multiplier, *κ* is the transition multiplier, and *π*_*j*_ is the stationary frequency of codon *j*. The standard codon model is therefore described by a 61 *×* 61 rate matrix. The diagonal entries of ***Q*** are chosen so that the rows sum to 0. Here, we consider Markov-modulated rate matrices, which allow for the rate matrix itself to change over the phylogeny (14). Specifically, in an *N* -category Markov-modulated rate matrix, we utilize a rate matrix of the form:

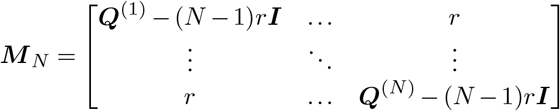

where each ***Q***^(*i*)^ is a whole codon rate matrix with a different *ω* parameter and, therefore, different *dN/dS*. All other parameters among the different ***Q*** matrices are shared. Additionally, we adopt the restriction that the *ω* parameter assigned to ***Q***^(*i*)^, denoted *ω*_*i*_, is given by

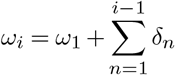

Where *δ* is just an increment from the last *ω* value. Therefore, each rate matrix going down the diagonal of ***M*** _*N*_ is strictly larger than the last. For each rate matrix ***Q***^(*i*)^, the stationary distribution is set to ***π****/N* so that within ***M*** _*N*_, the stationary distribution between selection regimes is uniform. Note that we only consider *r* to be a parameter of the model when *N >* 1.

Given a Markov-modulated rate matrix ***M*** _*N*_, we calculate the likelihood of an alignment using the transition probability matrix ***T*** = exp(***M*** _*N*_ *t*) and Felsenstein’s pruning algorithm (15).

### Dirichlet Process Prior

The Dirichlet process is a Bayesian nonparametric prior over distributions. That is, a draw

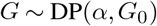

is itself a random probability distribution on a parameter space Θ, governed by a concentration parameter *α >* 0 and a base distribution *G*_0_, where supp(*G*_0_) = Θ. For any finite measurable partition (*A*_1_, …, *A*_*k*_) of Θ, the DP satisfies the defining property

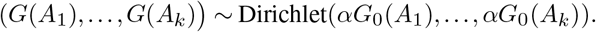

This characterization leads to a closed-form predictive distribution for sequential draws from *G*. If *θ*_1_, …, *θ*_*n*_ are i.i.d. from *G*, then the conditional distribution of *θ*_*n*+1_ given the first *n* values is

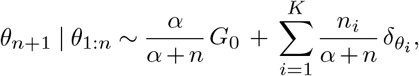

where *θ*_1_, …, *θ*_*K*_ denote the *K* unique values among the first *n* draws, *n*_*i*_ is the number of observations assigned to the *i*th unique value, and *δ*_*θ*_*i* is a Dirac-delta distribution with the location *θ*_*i*_.

The joint prior over the *K* unique parameter values and their associated counts can be written with *G* marginalized out:

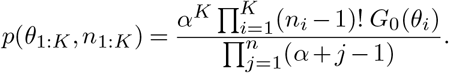

This expression depends only on the cluster sizes (*n*_1_, …, *n*_*K*_) and atom locations (*θ*_1_, …, *θ*_*K*_), and is invariant to permutations of the observation order. Its distribution over cluster assignments is described by the Chinese restaurant process (CRP), named after a clustering process imagined to take place in a Chinese restaurant (16). A customer enters a Chinese restaurant with an infinite number of tables and takes a seat at a table with a parameter value written on it. A second customer then enters and chooses a new table with probability *α/*(1 + *α*), or joins the first customer with probability 1*/*(1 + *α*). More generally, when the *j*th customer enters, they sit at a new table with probability *α/*(*j −*1 + *α*), or at an existing table *i*, with *n*_*i*_ customers already seated, with probability *n*_*i*_*/*(*j −*1 + *α*). The tendency for tables with many customers to attract additional customers is sometimes referred to as a “richget-richer” effect. When this prior is incorporated into a statistical model with a likelihood, each customer (data point) also has a “preference” for the parameter value associated with each table, in addition to preferring more crowded tables. The resulting posterior distribution naturally balances these parameter preferences against the tendency to cluster.

The CRP, therefore, provides a flexible prior for mixture models with an unknown number of components. In our model, we consider the prior

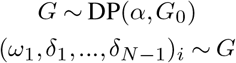

where (*ω*_1_, *δ*_1_, …, *δ*_*N−*1_) are the parameters defining our base non-synonymous/synonymous exchangeability rate (*ω*_1_) and the increments (*δ*_1:*N−*1_). This allows us to utilize an infinite site-wise mixture of Markov-modulated processes.

### Probabilistic Model

We consider two classes of models in this paper. The first model, which we will call the Markov-modulated codon model (CMM) resembles those originally investigated by (12). In our analyses, for a model with *N* submatrices in the Markov modulated rate matrix ***M*** _*N*_, we use the priors:

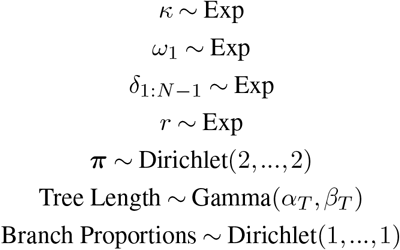

In our simulation analyses and globin demonstration, we set the exponential priors to have a rate of 3. In our simulations we set *α*_*T*_ and *β*_*T*_ such that E(Tree Length) = 35 and 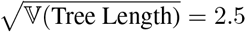. To express greater uncertainty in the true tree length, in our globin analysis we set *α*_*T*_ and *β*_*T*_ such that E(Tree Length) = 35 and 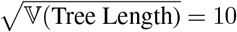. In the Dirichlet process mixture CMM (DPCMM), we allow each site to be assigned different values of *ω*_1_ and *δ*_1:*N−*1_. Therefore, the only difference in that model is the priors:

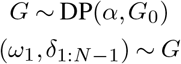

where *G*_0_ is the product of *N* Exp(3) distributions. The *α* parameter of the Dirichlet process is parameterized in terms of the expected number of partitions, which in all of our analyses is set to 3.

### Estimating Posterior Quantities

The objective of any Bayesian analysis is to estimate the distribution of the parameters given the data, *p*(*θ* |*X*), known as the posterior distribution. Except in special cases, this distribution does not have an analytically tractable form. Consequently, we rely on Monte Carlo methods to approximate posterior quantities. In this work, we use a Markov chain Monte Carlo (MCMC) sampler, which constructs a Markov chain whose stationary distribution is the posterior distribution of our model. Specifically, we employ the Metropolis–Hastings algorithm (17) to estimate parameter values, along with reversible-jump MCMC (RJ-MCMC) (18) to transition between parameter spaces corresponding to models with different numbers of evolutionary regimes.

The Metropolis-Hastings algorithm proceeds as follows:

#### Algorithm 1

Metropolis-Hastings

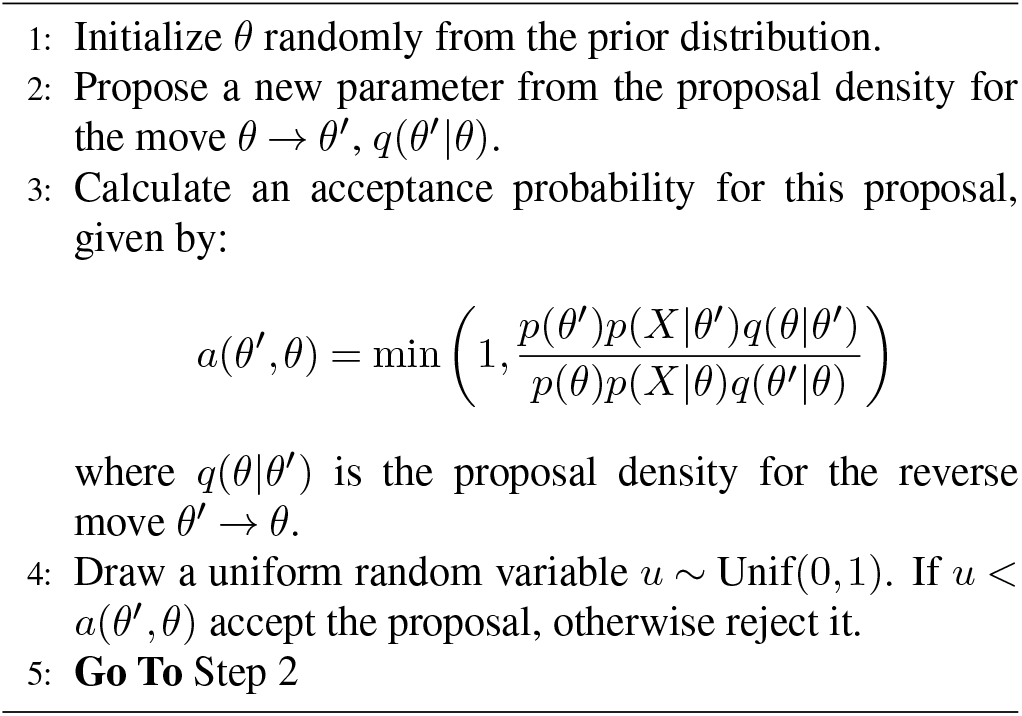

The fraction *q*(*θ*|*θ*^*’*^)*/q*(*θ*^*’*^|*θ*) is often called the Hastings ratio.

RJ-MCMC utilizes Metropolis–Hastings proposals that properly account for changes in the dimension of the posterior distribution, such as adding or removing a parameter. In the case of a pure birth/death RJ-MCMC move, a new parameter *θ*^*’*^ is drawn from a probability distribution *p*(*θ*^*’*^) or an existing parameter is deleted. The forward proposal density is given by

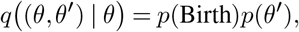

where *p*(Birth) is simply the probability that the MCMC kernel chooses to create a new parameter. The reverse proposal density, corresponding to the death move, is given by

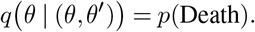

However, when the new parameter *θ*^*’*^ depends on the previous set of parameters *θ*, the Hastings ratio acquires an additional term to ensure reversibility. For example, consider a proposal *θ*_1_ *→* (*θ*_1_^*’*^, *θ*_2_^*’*^) that splits *θ*_1_ into two new parameters, or a proposal (*θ*_1_, *θ*_2_) *→ θ*_1_^*’*^ that merges two parameters into one. For the split move, we might draw *u ∼* Beta(5, 5) and define

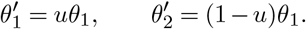

The corresponding merge move is defined by

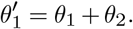

To ensure that the MCMC algorithm remains valid, RJMCMC requires including a determinant Jacobian factor in the acceptance probability (18). This is because the proposal couples the parameter space in a non-trivial way. The acceptance probability for the split move is

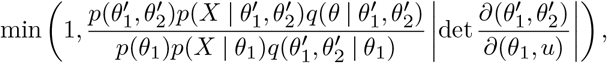

where

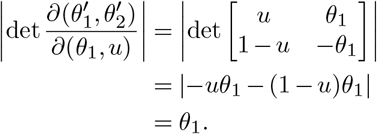

Because the merge move is the inverse of the split move, it uses the inverse of this determinant, 1*/θ*_1_^*’*^ .

We implemented a suite of MCMC moves within a Metropolis-within-Gibbs scheme, in which each iteration selects a single parameter type and applies several Metropolis–Hastings updates to that type. Dirichlet simplex moves, as implemented in RevBayes (19), were used to update *π* and branch proportions. Scaling moves were applied to update all scalar parameters (Tree Length, *κ, r, ω*_1_, *δ*_1:*N−*1_).

Exchange moves were used to update parameter values in (*ω*_1_, *δ*_1:*N−*1_) by transferring a proportion *p ∼* Unif(0, 1) of the mass from one selected parameter to another. We also developed a re-indexing move that swaps the values associated with two indices in (*ω*_1_, *δ*_1:*N−*1_). In both proposals, two indices are uniformly selected without replacement, and their corresponding parameter values are exchanged. Letting *i < j* denote the selected indices, under the cumulative parameterization of the category-specific *dN/dS* values, both of these moves affect only categories with indices in [*i, j*), while leaving all other categories unchanged.

Pure birth/death moves were added to increase/decrease the number of *δ* parameters and to de-instantiate/re-instantiate *r* when transitioning between *N* = 1 and *N* = 2; and split/merge moves were implemented to combine adjacent *δ*_1:*N−*1_ and *ω*_1_ parameters and split existing parameters into two (18, 20). Notice that in the case of the split-merge moves, the *dN/dS* values for all other indices besides the split/merged parameter remain unchanged due to the cumulative parameterization.

Neal’s algorithm 8 (21) was used to sample component assignments and parameter values from the Dirichlet process in the DPCMM model. The algorithm approximates the probability of assigning a site to a new CRP component using *m* auxiliary components, each with parameter values drawn from *G*_0_. It evaluates the likelihood of the site under each existing CRP component and each auxiliary component. The site is then assigned to one of these components according to its likelihood and prior probability, as in a Gibbs sampling update.

In our CMM model, we restricted the number of selection regimes to *N ≤* 5. At *N* = 5, the rate matrix ***M*** _5_ is a 305 *×* 305 rate matrix, and the likelihood evaluations become computationally intense. Although it is theoretically possible to search for even more regimes, it is computationally challenging in our implementation. In the DPCMM model, the split/merge and birth/death moves must operate on every cluster in the Dirichlet process, making these updates challenging. Additionally, increasing the number of selective regimes increases the dimension of the base distribution of the Dirichlet process, making Neal’s algorithm 8 less efficient. Additionally, we use eigen decomposition to compute our transition probabilities, meaning that the memory cost of the eigenvectors and eigenvalues for larger matrices, potentially at a per-site basis, becomes unreasonable. For this reason, we set a more conservative limit of *N ≤* 3. However, because the DPCMM model incorporates site heterogeneity in the evolutionary process, it is still potentially more sensitive than the CMM model in situations where fewer than 4 selection regimes are used at a given site, yet many more regimes exist across the full sequence.

We estimate the posterior *dN/dS* at every element of the MSA (each site in each taxon) using ancestral sampling. At each MCMC sampling iteration, we perform ancestral sampling in a pre-order traversal of the phylogeny, reconstructing the latent rate category for each node at each site. The *dN/dS* for latent rate category *k* belonging to ***Q***^(*k*)^ is computed as

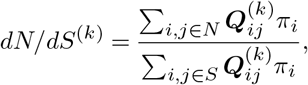

where *N* and *S* denote the indices of non-synonymous and synonymous transitions, respectively (22). Across the full MCMC run, this yields a joint posterior distribution of *dN/dS* over all taxa and all nodes of the tree.

Our MCMC implementation was validated by targeting the prior distributions and verifying that the resulting marginal posteriors matched the specified priors.

Our software outputs “trace” files from the MCMC sampler, which record the state of the chain at each sampled iteration. Additional posterior quantities of interest can be computed directly from these trace files using simple procedures. For example, one may wish to compute the posterior probability that two lineages, *A* and *B*, have *dN/dS* values (denoted *V* here) within some threshold *E* at a site *i*:

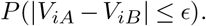

This probability can be estimated directly using the tip *dN/dS* trace file as a collection of posterior samples:

#### Algorithm 2

Compute Posterior Quantity

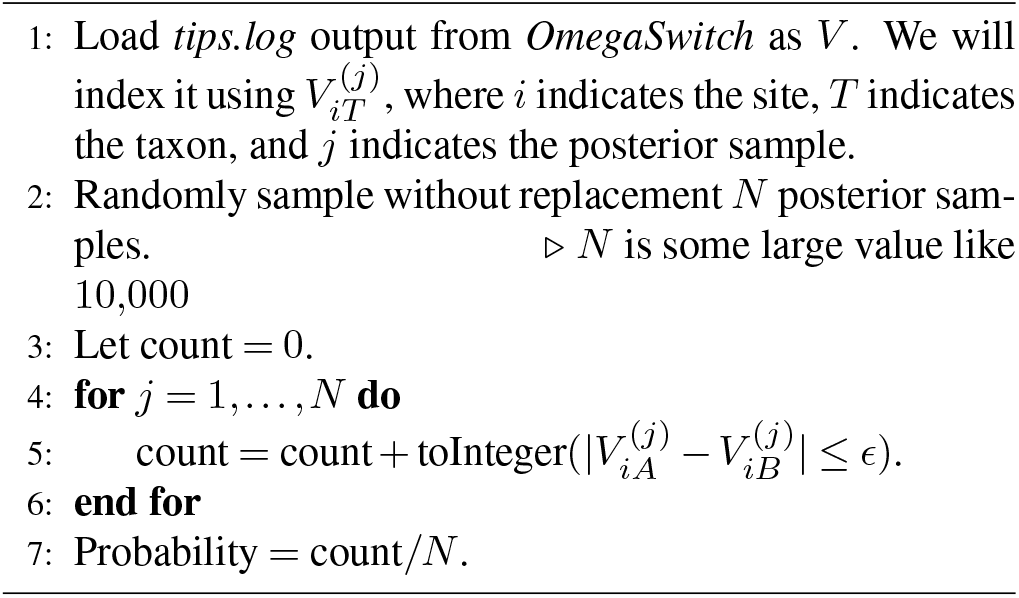

More generally, the logical expression in step 5 can be replaced by any quantity of interest that can be computed from the posterior samples. For example, a simple probability of positive selection at site *i* and taxon *A* might use

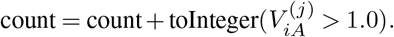

*OmegaSwitch* is also able to reconstruct ancestral *dN/dS* and is therefore capable of computing posterior quantities about how selection has changed over time. Please note that although this is standard in Bayesian inference, using an RJMCMC sampler introduces additional complications. Posterior summaries of quantities that are defined in every sampled model, such as *dN/dS*, can be computed in the usual way. However, summarizing parameters whose dimensionality differs across models (or is absent in some models), such as the *δ* parameters or *r*, requires additional care.

### Analyses

Simulation analyses were performed to assess the posterior coverage of two primary quantities of interest: the number of regimes in the Markov-modulated model and the *dN/dS* value at each site in the alignment. We simulated 50 alignments, each with 200 sites and 25 taxa, under the priors of the CMM and DPCMM models. Additionally, we ran the DPCMM model with the correct number of evolutionary regimes pre-specified to simulate (11). Each model was run with a burn-in of 5,000 iterations followed by 30,000 iterations, sampling every 100 iterations. We ran these simulations in parallel using the Zaratan HPC at the University of Maryland. Our initial experiments showed that these chain lengths yielded reasonable approximations to the posterior. Details of these analyses are recorded in **Table 2**.

**Table 1.** Posterior coverage frequencies for the 95% credible intervals or sets under the CMM and DPCMM models. Coverage for *dN/dS* was computed across site–taxon pairs within each simulated alignment. FB denotes the “Full Bayesian” DPCMM run in which the number of regimes and CRP components are jointly inferred, while FRC denotes the “Fixed Regime Count” run.

| Model | # Regimes | $dN/dS$ |
| --- | --- | --- |
| CMM | 1.00 | 0.96 |
| DPCMM (FB) | 1.00 | 0.96 |
| DPCMM (FRC) | – | 0.95 |

**Table 2.**
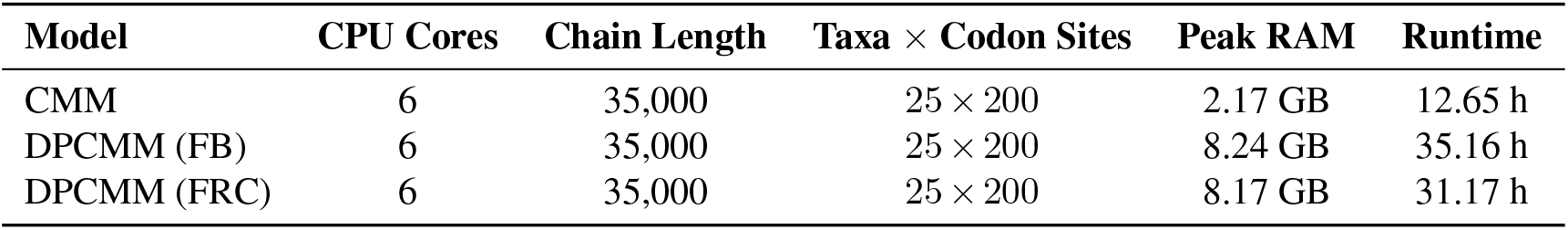
Computational requirements for the simulation datasets, including dataset size, chain length, core count, average peak memory usage, and average runtime. Simulations were run on the Zaratan HPC standard partition using dual AMD EPYC 7763 CPUs. FB denotes the “Full Bayesian” DPCMM run in which the number of regimes and CRP components are jointly inferred, while FRC denotes the “Fixed Regime Count” run.

To demonstrate the posterior quantities produced by *OmegaSwitch* on an empirical dataset, we analyzed vertebrate *α*- and *β*-globins. Globins provide a useful illustrative system because they are deeply conserved, have well-characterized structural constraints, and include paralogous proteins that have diversified across vertebrate evolution. Full-length*α*- and *β*-globin coding sequences were manually selected from GenBank. This alignment included *α*- and *β*- globins from *Homo sapiens* (NM_000517.6, CR541913.1), *Bos taurus* (LT548196.1, LT548198.1), *Gallus gallus*(NM_001004376.4, NM_205489.3) *Chordeiles minor* (KY485222.1, KY485233.1), *Anser indicus* (MH375701.1, MH375708.1), *Bombina bombina* (XM_053695065.1, XM_053695009.1), *Xenopus borealis* (M32453.1, M32456.1), *Chelonia mydas* (XM_037911157.2, XM_037907759.2), *Pelusios castaneus* (LC483570.1, LC497250.1), *Crocodylus niloticus* (MN905603.1, MN905616.1), *Danio rerio* (NM_131257.3, NM_131020.3), *Cyprinus carpio* (XM_042719705.1, XM_019117933.2), and *Salmo salar* (NM_001123662.1, NM_001123666.1). These taxa were selected to provide broad vertebrate representation while retaining complete coding sequences for both paralogs. Alignments were generated using the protein-coding sequences of the globins with MAFFT G-INS-i (23), and the codon-level alignments were generated with PAL2NAL (24). We fixed the known species tree for these organisms and analyzed the data under both CMM and DPCMM models. The samplers were run with a burn-in of 10,000 iterations, followed by an additional 50,000 iterations, sampling every 100 iterations in duplicate. Convergence of the chains was assessed using TRACER (25).

To analyze the relationship between the structure of the hemoglobin tetramer and *dN/dS*, the relative solvent availability (RSA) of the crystallographic structure of the tetramer (26) was computed using FreeSASA (27). Weighted contact number (WCN) of each *α*-carbon in the globin tetramer was computed using the MDanalysis package (28, 29).

Downstream analyses of the posterior distributions were done using a combination of R and Python scripts. Figures were generated using ggplot2 (30), and the visualization of *dN/dS* on the 3D structure of the hemoglobin tetramer was performed using ChimeraX (31).

## Results

Simulation analyses revealed that all versions of the models considered here have calibrated posteriors for the *dN/dS* values at each value in an MSA (**Table 1**). Likewise, the credibility sets for the number of evolutionary regimes also had calibrated coverage under our simulations. Additionally, on average, the true evolutionary regime had the largest posterior mass assigned to it (**Supplemental Material**).

In the globin demonstration, the CMM and DPCMM produced broadly similar posterior dN/dS estimates across site–taxon combinations, with differences likely reflecting the greater site-wise flexibility of the DPCMM. Notably, the DPCMM model had a higher inferred *r* parameter, which we discuss below.

When analyzing the relationship between relative solvent availability (RSA) in the globin tetramer and *dN/dS*, both models show a moderate correlation (Spearman correlation (CMM) = 0.274, Spearman correlation (DPCMM) = 0.269). The weighted contact number (WCN) showed a slightly weaker correlation with *dN/dS* (Spearman correlation (CMM) = -0.218, Spearman correlation (DPCMM) = -0.198).

## Discussion

### Empirical Demonstration

Globins are a diverse and ancient protein family, dating back to before the last universal common ancestor (32). In Metazoa, they are known for their diversity of secondary and tertiary structures (33), with the vertebrate hemoglobin tetramer being a central protein complex for oxygen delivery throughout the body. The human hemoglobin tetramer consists of two *αβ* heterodimers and, at full capacity, binds four heme molecules, which coordinate the storage of oxygen (34).

Our models recapitulate the expected features of hemoglobin, in particular, a low *dN/dS* in the heme-binding pocket (**Figure 1**). Upon inspecting the heme-binding pocket, we noticed interesting polarization across the *α*-helix holding the distal histidine. Specifically, a high *dN/dS* was observed on residues facing out towards the environment, while there was low *dN/dS* for the inward-facing residues of the helix. This pattern agrees with the literature on the relationship between *dN/dS* and protein structure. Early in this literature, an inverse relationship was observed between the solvent availability of a residue and amino acid substitution rates (3, 35–37). Later results showed that the weighted contact number for each residue may correlate more strongly (3, 38, 39). Both of these observations can be attributed to correlations of catalytic sites, which tend to be tucked away in the protein, and the stress induced by mutating a site, which is often more severe in densely packed regions of a protein’s structure (3). Indeed, globin proteins are a particularly interesting case to look at for historical reasons, as literature dating back to the 1960s has discussed the differences in conservation between the interior and exterior of the protein (40).

**Fig 1.**
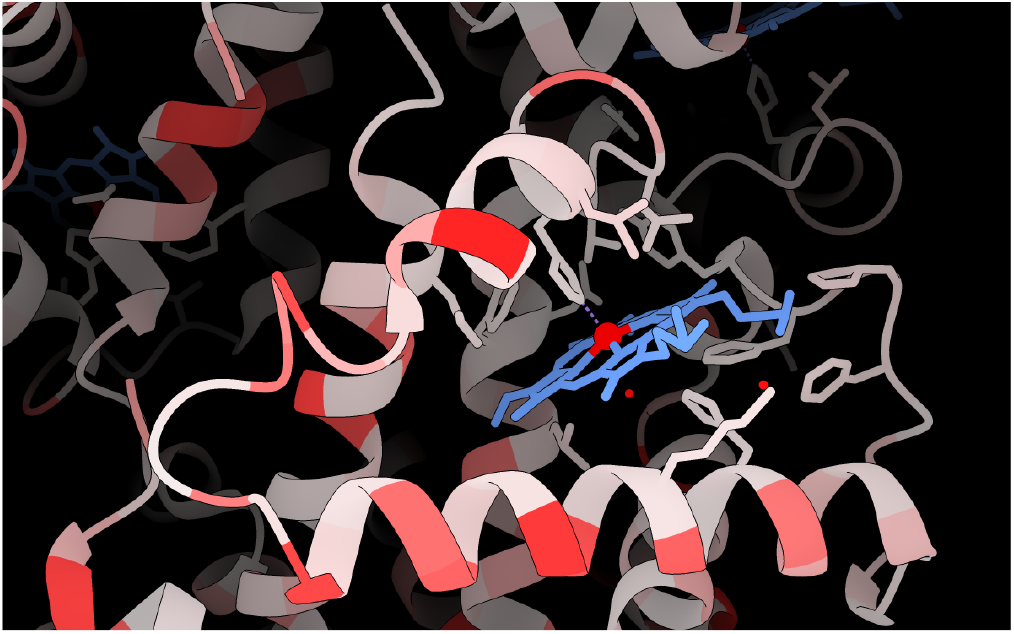
Heme bonding pocket of a human *α*-globin subunit in the globin tetramer. White indicates low median posterior *dN/dS* and red indicates high median posterior *dN/dS*, with a maximum of 3.5. The heme molecule is colored blue, with the iron atom in the middle colored red. Notice the conservation on the inside of the heme pocket and polarized selection across the *α*-helix below, with greater *dN/dS* being assigned to solvent-exposed parts of the protein.

To investigate this in the case of *α*- and *β*-globin, we computed the relative solvent availability (RSA) and weighted contact number (WCN) for each residue in the hemoglobin tetramer. There was a mild correlation between RSA and *dN/dS* and a slightly weaker correlation between WCN and *dN/dS* (**Figure 2**). Although this is the opposite of what has been reported for these two metrics in the past (WCN usually shows a stronger correlation than RSA), we are studying a single protein family rather than a large multi-gene dataset. Additionally, prior findings used different models to estimate *dN/dS* than ours; thus, their estimates may differ from the posterior median *dN/dS* reported here (35, 37). Other studies use site-specific evolutionary rates that don’t strictly correspond to *dN/dS* estimates (39). These results illustrate how *OmegaSwitch* posterior summaries can be integrated with structural information to examine variation in selective constraint across sites and lineages.

**Fig 2.**
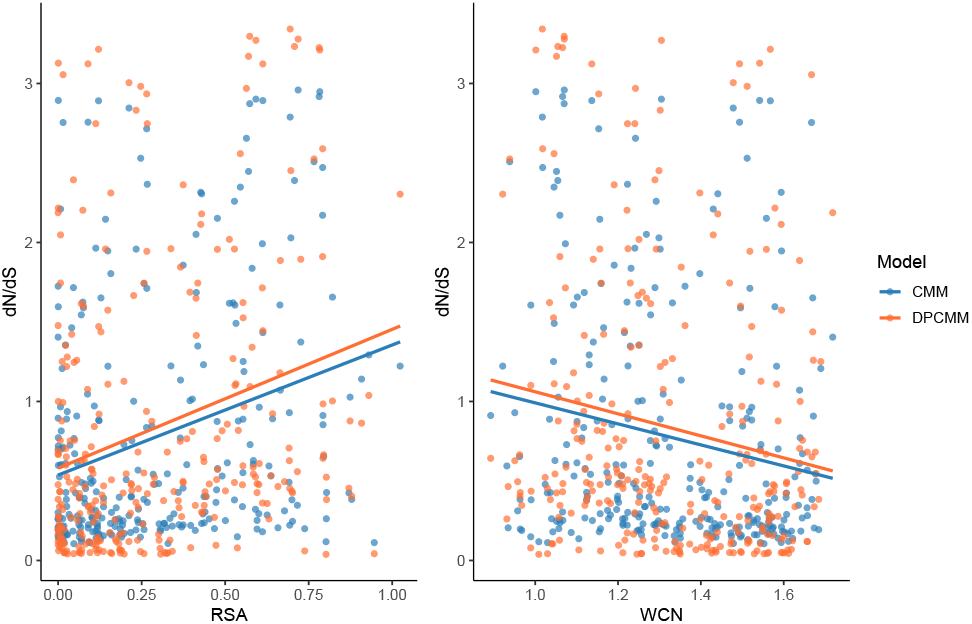
Relationship between relative solvent availability (RSA), Weighted Contact Number (WCN), and *dN/dS* estimates for the human globin tetramer.

Although there is general agreement among the models on the posterior *dN/dS* per site in human globins (**Figure 3**), there are also differences in the posterior median and variance. This is likely partially explainable by the low transition rate *r* between Markov-modulated categories. On average, this parameter was approximately 0.06 in the CMM model and approximately 0.26 in the DPCMM model (**Figure 4**). That approximately 4-fold difference in the transition rates between *dN/dS* categories likely explains why the DPCMM model is freer to infer larger *dN/dS* differences. A possible reason is that the DPCMM model is less affected by sites unlikely to have changed over evolutionary time. These sites would keep the transition rate low in the CMM model, whereas in the DPCMM model, they may be assigned to a category with relatively similar rates across all *dN/dS* classes. Consider, for example, site 1 in this model, which does not deviate from methionine. When we look at the posterior distribution of *ω* under the DPCMM model, it looks like the first two classes of the CMM model (**Figure 5**). On the other hand, site 116 has a posterior distribution with *ω* parameters similar to the full CMM classes 1, 2, and 5 (**Figure 5**). Therefore, in the DPCMM model, each site is allowed to occupy its preferred class across the tree without suppressing the transition rate between categories for other sites.

**Fig 3.**
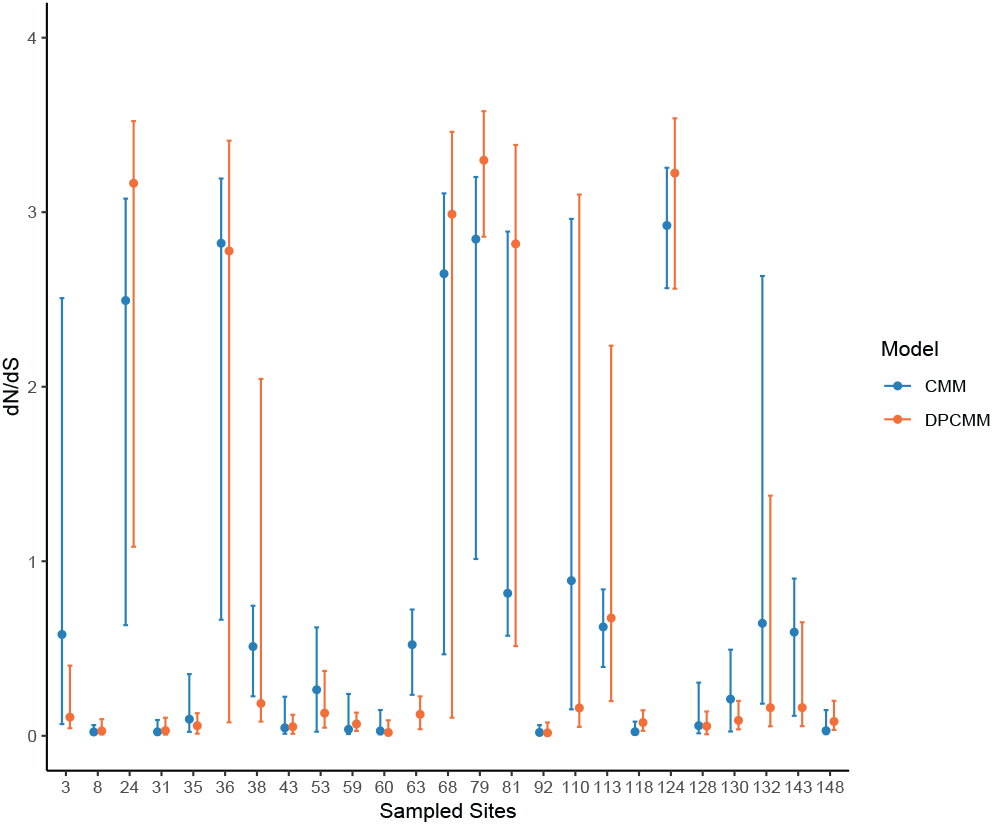
Comparison of the posterior median and inter-quartile range for *dN/dS* of randomly selected sites in human *α*-globin inferred by the CMM and DPCMM model.

**Fig 4.**
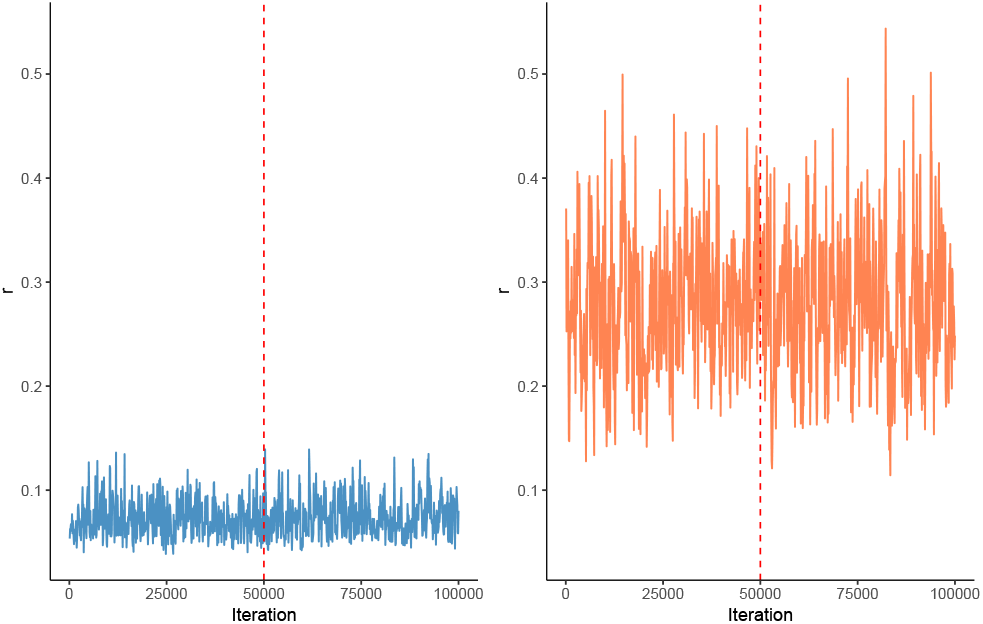
Comparison of the *r* parameter (the rate of swapping between evolutionary regimes) concatenated trace in the CMM (left) and DPCMM (right) model. The dashed line indicates where the first MCMC chain ends and the second chain begins.

**Fig 5.**
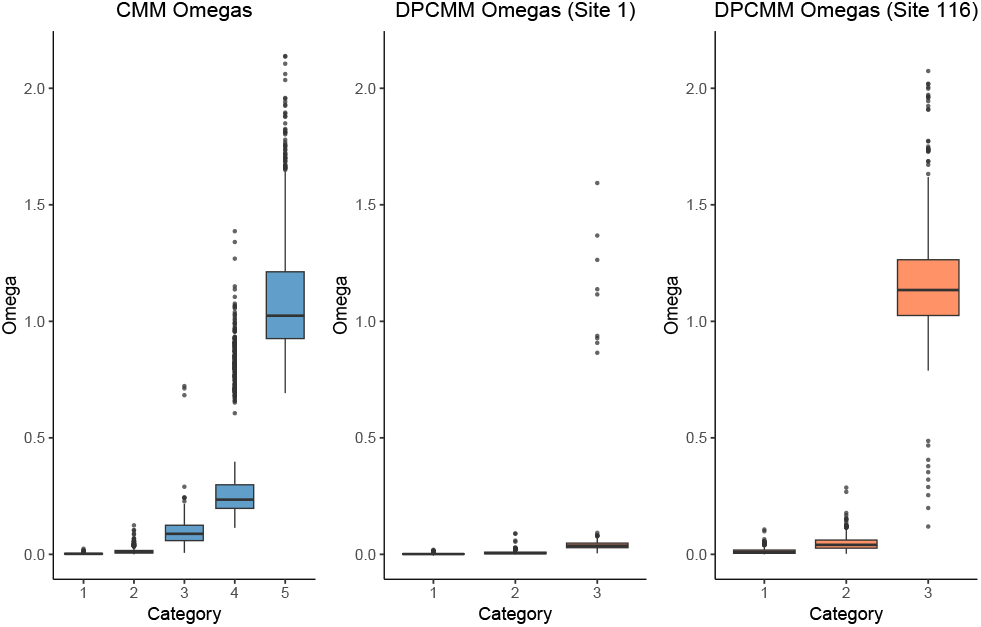
Comparison of the posterior distribution of *ω* parameters assigned by the CMM model and DPCMM model. The CMM plot corresponds to the global posterior, while the DPCMM plots are posteriors for particular sites in the alignment.

To demonstrate the flexibility of the Bayesian framework for investigating biological questions about protein evolution, we used posterior samples from *OmegaSwitch* to calculate the probability that human *α*-globin has a greater *dN/dS* than human *β*-globin at each homologous site (**Figure 6**). Because *OmegaSwitch* produces samples from the joint posterior distribution of site- and lineage-specific *dN/dS*, such quantities can be calculated directly after inference without defining an additional statistical model or hypothesis test. More generally, posterior samples can be transformed to address arbitrary comparisons among sites and lineages.

**Fig 6.**
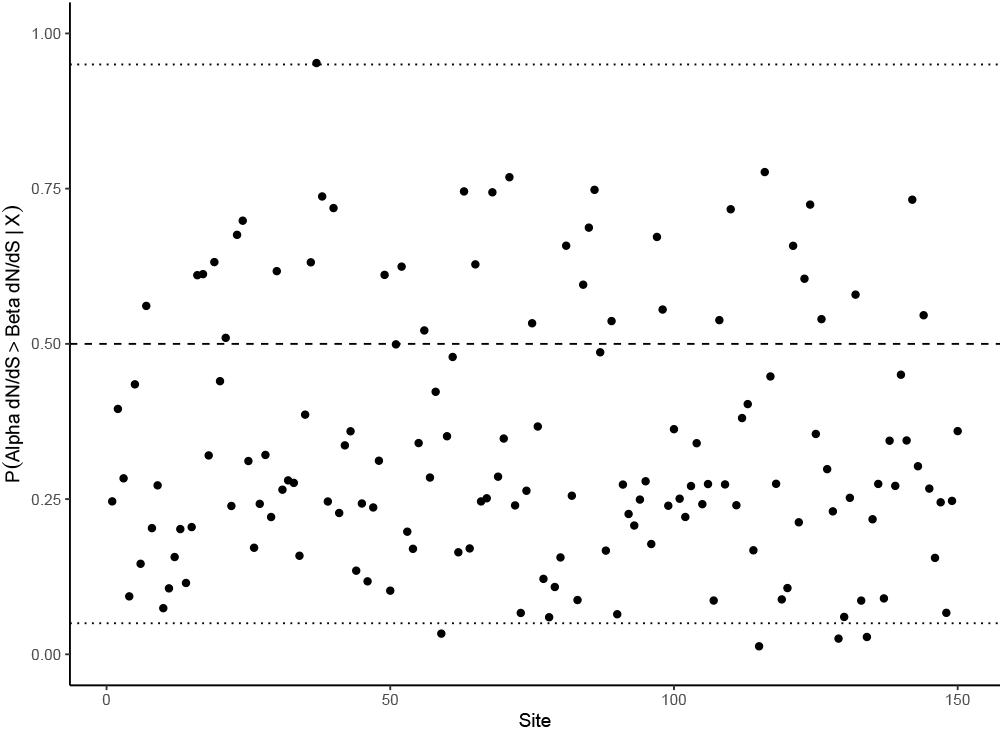
CMM posterior probabilities of the *dN/dS* of human *α*-globin being greater than that of human *β*-globin. The horizontal lines indicate 0.05, 0.5, and 0.95 posterior probability, respectively.

One site was particularly notable, with approximately 0.95 posterior probability of greater *dN/dS* in *α*-globin than in *β*- globin. This probability should be interpreted as a directional posterior probability and not confused with a hypothesis-testing threshold. This comparison corresponds to *α*35 and *β*34, residues near the C-terminal end of the B helix that contribute to the *α*_1_*β*_1_ dimer interface. *β*34 is a conserved Val whose substitution perturbs the *α*_1_*β*_1_ contact region (41), whereas *α*35 is a Ser that makes inter-subunit contacts within the same interface (42, 43). Consistent with the inferred difference in evolutionary constraint, documented substitutions at these positions appear to have substantially different structural and functional consequences. Reported variants of *β*34 result in increased oxygen affinity in every variant recorded in the HbVar database (44), thereby reducing oxygen delivery to peripheral tissues. Notably, these variants include substitutions of several chemically distinct residues, such as Phe, Leu, Ala, and Asp; this suggests *β*34 is a strongly biochemically constrained residue. *α*35 also has reported variants in HbVar associated with unstable tetramers (44), but the documented variants at this position involve substitution to bulky residues (Pro, Phe, and Tyr). Consistent with this apparent difference in constraint, *α*35 exhibits greater amino-acid variation across the vertebrate alignment, occurring as Ala, Val, Thr, His, or Ser. In contrast, *β*34 is conserved as Val in all sampled taxa except *X. borealis*, in which it is Thr.

### Conclusion

Our software *OmegaSwitch* provides a documented user interface and enables analyses such as those presented here. Although we have highlighted only a subset of the posterior quantities produced by the model, the framework naturally supports many additional summaries. By placing a distribution over the parameters governing the data-generating process, the model yields a wide range of internally consistent posterior quantities. Beyond introducing DPCMM, *OmegaSwitch* provides an accessible C++ implementation of foundational work of (12) and site-heterogeneous Dirichlet process (11) models. We anticipate that this software will be broadly useful for investigating evolutionary selection both between homologs and across sites.

## Data Availability Statement

The software used to perform these analyses was written in C++ and designed for use in the terminal. *OmegaSwitch* and compilation/usage instructions are available at https://github.com/Wesley-DeMontigny/OmegaSwitch/tree/main. All additional files related to the analyses performed will be made public after peer review.

## ACKNOWLEDGEMENTS

The authors acknowledge the University of Maryland (UMD) supercomputing resources (https://hpcc.umd.edu) made available for conducting the research reported in this paper. WCD was partially funded through a TAship from UMD and by the Gordon and Betty Moore Foundation through Grant GBMF11481 (https://doi.org/10.37807/GBMF11481) to UMD. The authors thank John Huelsenbeck for his guidance and help getting started with phylogenetic modeling and software development.

